# Analysis and visualization of quantitative proteomics data using FragPipe-Analyst

**DOI:** 10.1101/2024.03.05.583643

**Authors:** Yi Hsiao, Haijian Zhang, Ginny Xiaohe Li, Yamei Deng, Fengchao Yu, Hossein Valipour Kahrood, Joel R. Steele, Ralf B. Schittenhelm, Alexey I. Nesvizhskii

## Abstract

The FragPipe computational proteomics platform is gaining widespread popularity among the proteomics research community because of its fast processing speed and user-friendly graphical interface. Although FragPipe produces well-formatted output tables that are ready for analysis, there is still a need for an easy-to-use and user-friendly downstream statistical analysis and visualization tool. FragPipe-Analyst addresses this need by providing an R shiny web server to assist FragPipe users in conducting downstream analyses of the resulting quantitative proteomics data. It supports major quantification workflows including label-free quantification, tandem mass tags, and data-independent acquisition. FragPipe-Analyst offers a range of useful functionalities, such as various missing value imputation options, data quality control, unsupervised clustering, differential expression (DE) analysis using Limma, and gene ontology and pathway enrichment analysis using Enrichr. To support advanced analysis and customized visualizations, we also developed FragPipeAnalystR, an R package encompassing all FragPipe-Analyst functionalities that is extended to support site-specific analysis of post-translational modifications (PTMs). FragPipe-Analyst and FragPipeAnalystR are both open-source and freely available.

## INTRODUCTION

The FragPipe computational platform (https://fragpipe.nesvilab.org/) is increasingly popular in the proteomics field for processing proteomics datasets using different quantification strategies, including data-dependent acquisition (DDA) label-free quantification (LFQ)^1^, tandem mass tag (TMT)^2^, and data-independent acquisition (DIA) LFQ^3, 4^. While FragPipe produces well-formatted output tables that are ready for analysis, further downstream statistical analyses and visualizations are typically required to interpret the results. Therefore, there is an unmet need for a user-friendly downstream statistical analysis and visualization tool that can support advanced bioinformatics analyses. Reproducibility remains an unsolved issue in the scientific research community.

In the field of quantitative proteomics, a range of software packages has been developed over time to aid scientists in analyzing their data, including DEP^5^, protti^6^, Perseus^7^, LFQ-Analyst^8^, and MSstats^9, 10^. However, most of them focus on the MaxQuant^11^ output and often only consider LFQ datasets^5, 6, 8, 12^. Few support TMT and DIA datasets and/or require users to have programming proficiency^6, 10^. Furthermore, the majority of tools only use quantitative data summarized to the protein level as the input, even though peptide-level quantification may offer more accurate biological interpretation of the data in certain applications^13^.

To support the growing body of FragPipe users and the diversity of proteomics data analysis workflows offered by FragPipe, we created FragPipe-Analyst, an R Shiny website. FragPipe-Analyst is based on the previously described LFQ-Analyst^8^ code, which we extended to support the diverse outputs covering all major quantification workflows (LFQ, TMT, and DIA) from the various FragPipe workflows, along with various feature improvements in the analysis and visualization steps. FragPipe-Analyst offers a range of useful functionalities, such as multiple missing value imputation options, normalization, data quality control, unsupervised clustering, differential expression analysis using Limma^14^, and gene ontology (GO)/pathway enrichment analysis using Enrichr^15^ to bridge the gap between proteomic search results and downstream analysis. Moreover, we developed FragPipeAnalystR, an R package encompassing all FragPipe-Analyst core functionalities and additional site-specific analyses for post-translational modification (PTMs) data.

## METHODS

### Design and Implementation

The majority of functionalities available in FragPipe-Analyst were described in the LFQ-Analyst manuscript^8^. However, FragPipe-Analyst has been substantially extended to include more interactive features and increased flexibility. We also created a standalone R package, FragPipeAnalystR, to support a broader community with more advanced features. FragPipe-Analyst and FragPipe-AnalystR share the same common modular design with the following modules: (1) I/O module that handles the input and output files. (2) Data manipulation module which has functions for operating the fundamental data structure, including remove/merge samples and feature selections. (3) Data filtering module which provides methods for filtering data based on missing values. (4) Normalization module that provides several normalization methods. (5) Imputation module that provides several imputation methods. (6) Quality control (QC) module, which provides functions to generate various visualizations, including principal component analysis (PCA) plots and heatmaps. (7) Differential expression (DE) analysis module, which provides functions for performing statistical procedures and result visualization, such as a volcano plot. (8) Enrichment analysis module which provides statistical procedures specifically for inferring biological insights, such as overrepresentation tests.

In terms of data processing, FragPipe-Analyst and FragPipe-AnalystR first take input files (Table 1) produced by FragPipe and perform preprocessing steps to remove contaminants (proteins or peptides), transform the intensity values into log2 scale (except for spectral count data in LFQ), and harmonize them into the Bioconductor ‘SummarizedExperiment’ object. Internally, FragPipe-Analyst and FragPipe-AnalystR adopt the SummarizedExperiment as the main data structure, and the data object is further processed by different modules. Optional missing value filtering, data normalization, and missing value imputation can also be applied.

**Table 1.** Input files for FragPipe-Analyst based on quantification workflows used in FragPipe.

|  | DDA LFQ | TMT | DIA LFQ |
| --- | --- | --- | --- |
| Quantification<br>Report | combined_protein.tsv<br>or<br>combined_peptides.tsv | abundance/ratio_gene report,<br>abundance/ratio_protein report,<br>or abundance/ratio_peptide<br>report (.tsv) | protein group matrix<br>(.pg_matrix) or<br>precursor group<br>matrix (.pr_matrix) |
| Experiment<br>Annotation | experiment_annotation.tsv |  |  |

As the first step of data processing in FragPipe-Analyst, input quantification data can be optionally filtered to remove entries with too many missing values (controlled by ‘Min percentage of non-missing values globally’ and ‘Min percentage of non-missing values in at least one condition’). When reading DDA LFQ data generated by IonQuant^1^, the user can specify Intensity, MaxLFQ intensity, or Spectral Counts as quantitative measurements. By default, the input data are not normalized, as they are assumed to have already been normalized by the FragPipe quantification tools. However, the variance-stabilizing normalization option (performed using the R package vsn^16^) is available and may sometimes be beneficial for DDA-and DIA-based LFQ data. In the missing value imputation step, a missing not at random method, which uses random draws from a Gaussian distribution left-shifted by 1.8 standard deviation with a width of 0.3 (Perseus style imputation) is applied by default for DDA and DIA LFQ data. No imputation is performed for TMT data by default. Other imputation options are available, including k-nearest neighbor and MLE, as provided by the R package MSnbase^17^. The resulting table after filtering for missing values, data normalization, and missing value imputation is subsequently subjected to differential expression analysis and enrichment analysis.

In the differential expression analysis module, feature-wise (protein or peptide) linear models combined with empirical Bayesian statistics from Limma^14^ are used to test for statistically significant differences in abundance between conditions. Both p-values and adjusted p-values (by default, using the Benjamini-Hochberg method) are computed. For the enrichment analysis, the API provided by Enrichr^15^ is used, and hypergeometric tests are performed to determine the overrepresented categories. Enrichment analysis can also be performed using pathway databases (KEGG, Hallmark, or Reactome) or Gene Ontology (GO). Notably, we provide options for using either the entire proteome or the list of proteins identified in the experiment as the background for the enrichment analysis. Additionally, FragPipe-AnalystR supports gene set enrichment analysis using clusterProfiler^18^ as well as file conversion and result visualization for site-specific PTM analysis. Visualization from different modules relies mainly on the R packages ggplot2^19^, ComplexHeatmap^20^, and plotly^21^. The FragPipe-Analyst web interface was implemented as a Shiny app, as previously described^8^. The differential expression and enrichment analysis results and images (e.g., PCA, heatmaps, volcano plots) can also be downloaded for subsequent use in other computational tools and for publication purposes.

### Datasets

Several previously published datasets were downloaded and reanalyzed in this study to demonstrate FragPipe-Analysts capabilities: (i) an LFQ affinity purification mass spectrometry (AP-MS) interactome dataset, (ii) clear cell renal cell carcinoma (ccRCC) TMT global proteome and phosphoproteomics datasets, (iii) a ccRCC DIA whole proteome dataset, and (iv) a limited proteolysis coupled to mass spectrometry (LiP-MS) DIA dataset. In all cases, raw mass spectrometry files were first converted into an open mzML format using the msconvert utility of the Proteowizard software suite before further processing. All datasets were analyzed using the FragPipe computational platform (v20.0, https://fragpipe.nesvilab.org) using the settings described below.

*<u>LFQ AP-MS</u>* dataset: Raw files were downloaded from the PRIDE database (https://www.ebi.ac.uk/pride/) via the ProteomeXchange repository under the PXD019469 identifier. Seven raw files in total were downloaded (qx000121, qx001223, qx001225, qx001226, qx001227, qx001283, and qx001487) corresponding to three replicates of the CCND1 bait purification and four negative controls. The built-in FragPipe ‘LFQ-MBR’ workflow was used. Briefly, MSFragger^22^ (v3.8) was used to perform a closed search followed by label-free quantification with match-between-runs enabled using IonQuant^1^ (v1.9.8). In the closed search, both the initial precursor and fragment mass tolerances were set to 20 ppm. After mass calibration^23^, narrower tolerances were automatically selected by MSFragger. The enzyme was set to strict trypsin, and the maximum allowed missed cleavage was set to 2. Methionine oxidation and protein N-terminal acetylation were set as variable modifications, and carbamidomethylation of cysteine was set as a fixed variable modification. After the closed search, MSBooster^24^, Percolator^25^, and Philosopher^26^ were used to calculate the deep-learning scores, perform rescoring, and estimate the FDR for the peptide to spectrum matches (PSMs). In the label-free quantification stage, IonQuant^1^ was used and default settings were applied. Briefly, the mass tolerance was set to 10 ppm and the retention time tolerance was set to 0.4 minute. Match-between-run and MaxLFQ intensity calculations were enabled. The “min scans” parameter was set to 3, “min isotopes” was set to 2, and “MaxLFQ min ions” was set to 2. In addition to the analysis via FragPipe-Analyst presented here, the reprint.int.tsv file generated by FragPipe was submitted to the Resource for Evaluation of Protein Interaction Network (https://reprint-apms.org/) website to perform SAINTexpress^27^ analysis with default settings for comparison.

*<u>ccRCC TMT</u>* dataset: A subset of the samples from the global proteome and phosphoproteome dataset (four TMT 10-plexes out of twenty-three 10-plexes in total) was downloaded from Proteomic Data Common (PDC, https://pdc.cancer.gov/pdc/) under study PDC000127 and PDC000128, respectively. The subset included 36 samples, of which 32 were ccRCC patient samples (20 tumors, 12 normal tissues), three were technical replicates of the same QC sample, and one sample was determined to be a non-ccRCC tumor and was excluded from the analysis in the original study. The FragPipe default ‘TMT10-bridge’ and ‘TMT10-bridge-phospho’ workflows were used for global proteome and phosphoproteome data quantification, respectively. A common ccRCC pool sample was used as a bridge to quantify across multiple sets. Data were processed as described in the original publication^28^. MS/MS spectra were searched against a reviewed *H. sapiens* subset of the UniProt^29^ database and common contaminant sequences (downloaded on September 8, 2022, UP000005640, 20432 proteins) appended with an equal number of decoys. Whole cell lysate MS/MS spectra were searched using a precursor-ion mass tolerance of 20 ppm, allowing C12/C13 isotope errors of −1/0/1/2/3. Mass calibration and parameter optimization were performed. Cysteine carbamidomethylation (+57.0215) and lysine TMT labeling (+229.1629) were specified as fixed modifications, and methionine oxidation (+15.9949), N-terminal protein acetylation (+42.0106), and TMT labeling of peptide N-terminus and serine residues were specified as variable modifications. For the analysis of phosphopeptide-enriched data, the set of variable modifications also included the phosphorylation (+79.9663) of serine, threonine, and tyrosine residues. The search was restricted to tryptic peptides, allowing for up to two missed cleavage sites. Peptide-to-spectrum matches were further processed using Percolator^25^, converted to pep.xml format, and with the phosphopeptide-enriched dataset, pep.xml files were additionally processed using PTMProphet^30^ to localize the phosphorylation sites. The resulting pep.xml files were then processed using ProteinProphet^31^ and filtered to a 1% false discovery rate at the protein and PSM levels using the Philosopher toolkit^26^ v4.8. TMT quantification was extracted from MS/MS spectra using Philosopher, and the PSM output files were then further processed using TMT-Integrator v3.2.0, to generate summary reports at the gene, peptide, and modification site levels. Protein abundances were log2 transformed and median centered.

*<u>ccRCC DIA</u>* <u>dataset</u>: Raw files from the same set of 32 ccRCC samples as in the ccRCC TMT dataset (QC or the non-ccRCC tumor samples were not profiled using DIA) were downloaded from the PDC under study PDC000200. The FragPipe built-in ‘DIA_SpecLib_Quant’ workflow was used. In brief, MSFragger-DIA^4^ was used to perform database searching. The initial precursor and fragment mass tolerances were set at 20 ppm. After mass calibration, MSFragger-DIA adjusts the tolerance adaptively. Strict trypsin was selected as the digestion enzyme and the maximum allowed missed cleavage was set to 1. After the MSFragger-DIA search, MSBooster^24^ was used to calculate the deep-learning-based scores, and Percolator^25^ was used to perform the rescoring. Then, Philosopher was used to estimate the FDR and generate the intermediate report used by EasyPQP^3^ to build a spectral library. The library was filtered to obtain a 1% FDR at the protein and peptide levels. The DIA-NN^42^ quantification module was used to perform targeted quantification based on the spectral library and the original DIA spectra.

*<u>LiP-MS</u>* dataset: Raw files were downloaded from PRIDE via the ProteomeXchange repository using the PXD035183 identifier as described above. The data was processed using FragPipe ‘DIA_SpecLib_Quant’ workflow as described above, except enzyme specificity was changed to ‘semi-tryptic’ option.

## RESULTS AND DISCUSSION

### Overview of the FragPipe-Analyst

An overview of FragPipe-Analyst is shown in Figure 1. FragPipe-Analyst was designed for experiments containing samples with more than one condition. Each condition should have at least two samples. We adopted a typical workflow that consists of sequential normalization, filtering, and imputation steps, as previously described^5^ and focused on quality control and differential expression analysis. To use FragPipe-Analyst, users must upload two files: a quantification report and an experiment annotation file (experiment_annotation.tsv). Both files are obtained by executing FragPipe. The quantification report is the output based on the user’s choice of workflow (**Table 1**). The experimental annotation file contains information about the experimental design. It is automatically generated based on the user’s settings in FragPipe and may need to be manually checked and edited prior to its upload to FragPipe-Analyst. After ingesting these two files, FragPipe-Analyst combines them into an internal data structure. Further visualizations, including the QC and DE results, are then generated automatically.

**Figure 1.**
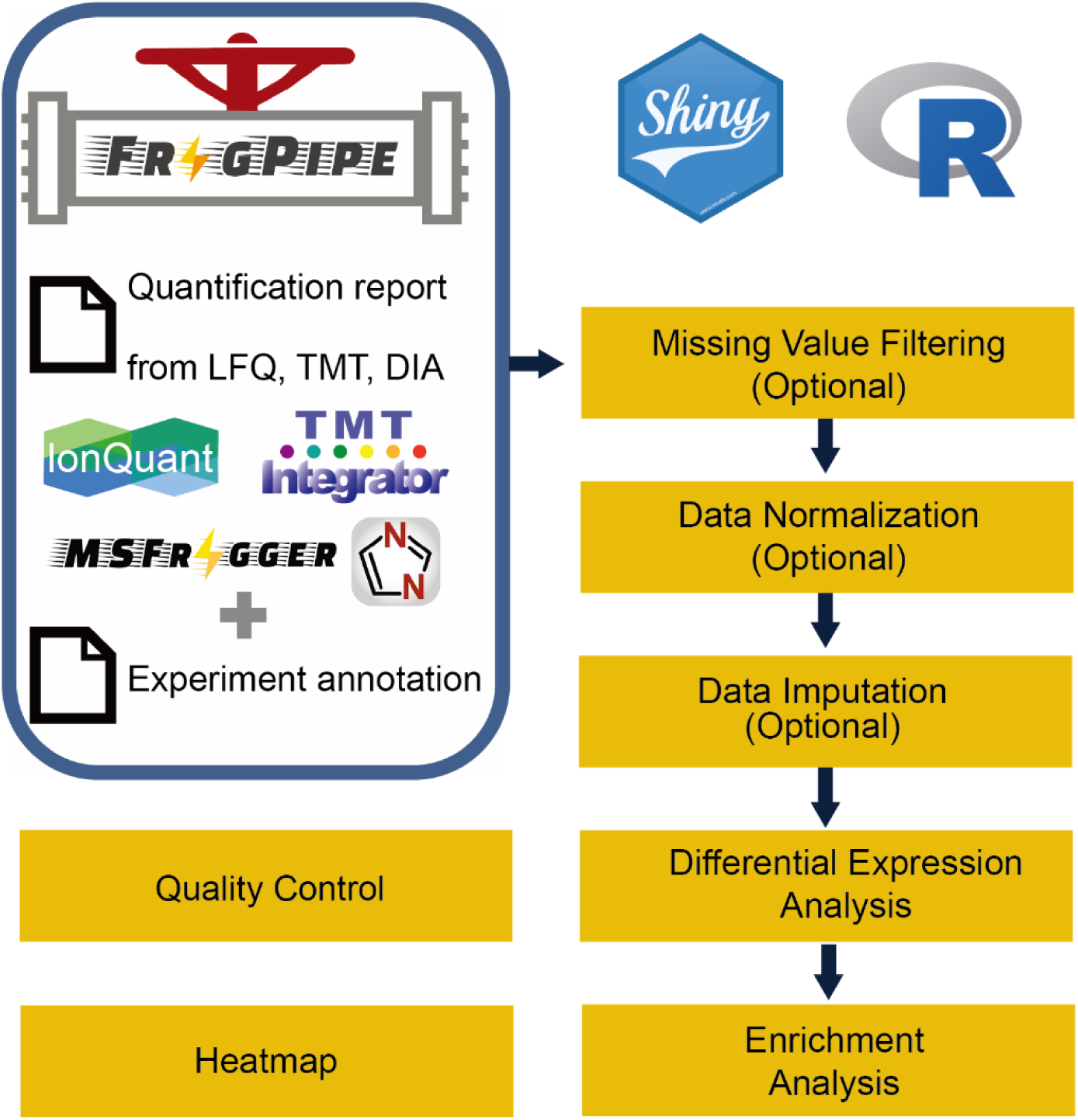
Overview of FragPipe-Analyst. LFQ: data-dependent label-free quantification, TMT: tandem mass tag, DIA: data-independent acquisition LFQ.

To illustrate the power and ease of use of FragPipe-Analyst, we describe how it can be used to re-analyze several previously published datasets, largely reproducing the original findings.

### Analysis of a TMT-based ccRCC cancer proteomics data

We first utilized FragPipe-Analyst to interrogate a global TMT-based proteomics dataset^28^ profiling clear cell renal cell carcinoma (ccRCC; the major kidney cancer subtype) patient samples. We only selected a subset of the data, four TMT-10 plexes out of 23 plexes in total for the demonstration. Note that although the original study was intended to profile ccRCC patients only, some samples were found to be of different kidney cancer subtypes such as papillary renal cell carcinoma (pRCC) and chromophore renal cell carcinoma. Data were processed using FragPipe^4^ under settings closely matching those used in the original study^28^ (see **Methods**). After finishing data processing in FragPipe and annotating samples, the quantification report and the experiment annotation files (abundance_protein_MD.tsv and experimental_annotation.tsv) were uploaded and analyzed by FragPipe-Analyst to quickly obtain results ready for interpretation (**Figure 2**). As shown in **Figure 2a**, a PCA plot implemented in plotly^21^ shows that all quality control (QC) samples from the different TMT-10 plexes formed a tight cluster. This indicates an overall high quality of quantification data (low technical variability). A clear separation between tumor (T) and normal adjacent tumor (NAT) samples from ccRCC patients was observed, underlining the biological differences between these two conditions. When using the dropdown menu to switch the PCA plot to be colored by plex number, which is a useful visualization option to assess batch effects (see **Supplementary figure S1 and 2** for additional QC plots), a clear batch effect is visible, suggesting that one of the plexes is notably different. Indeed, from the original study, we know that this is due to several plexes containing only tumor samples (imbalanced design). In the original manuscript, the batch effect stemming from unbalanced design was corrected using ComBat^32^. Here, we simply point out that our visualization tools assist users with the detection of such batch effects, so that an informed decision can be made regarding the need for batch correction.

**Figure 2.**
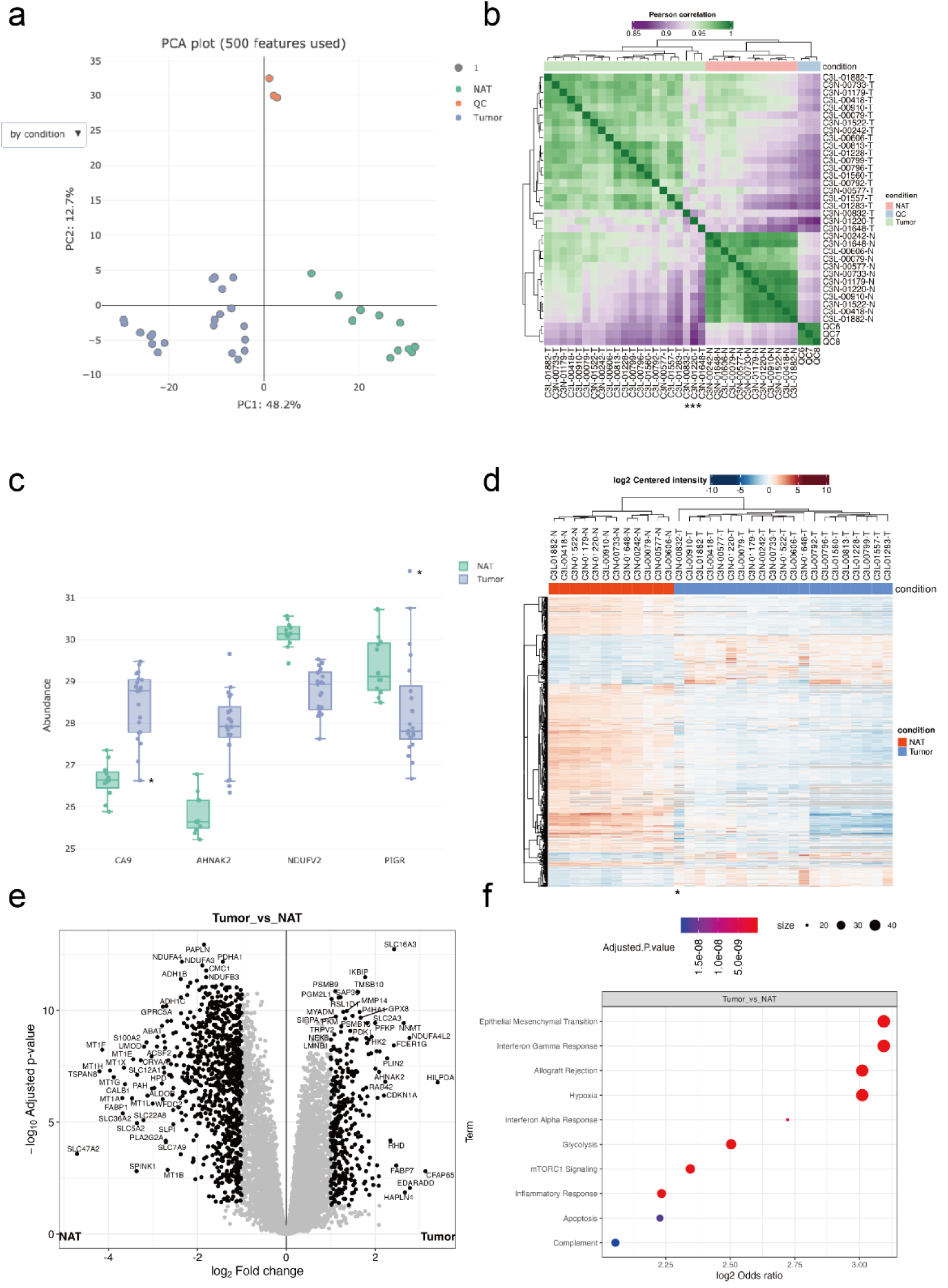
Example analysis results of global proteome ccRCC dataset obtained from FragPipe-Analyst. **(a)** PCA plot **(b)** Correlation matrix plot. Asterisks denote samples of dissimilar patterns. **(c)** Boxplot shows selected markers of renal cell carcinomas. Asterisks denote the expression of the papillary renal cell carcinoma sample (C3N-00832-T) **(d)** Heatmap based on differentially expressed proteins. Asterisk denotes the sample with dissimilar pattern. **(e)** Volcano plot **(f)** Over-representation test result for upregulated proteins identified in **(e)** against Hallmark gene set. Only proteins with log2 fold change more than 0.7 and adjusted p value lower than 0.05 were considered.

Correlation heatmaps are useful for QC and detecting outliers. Here, the correlation heatmap generated by FragPipe-Analyst (**Figure 2b**) aligned with our earlier observations from the PCA plot in such that tumors, NATs, and QC samples formed their own clusters. Interestingly, three tumors (C3N-00832-T, C3N-01220-T, and C3N-01648-T) showed dissimilar patterns. From the user’s perspective, knowing that renal cell carcinomas (RCCs) other than ccRCC may unintentionally be misclassified at the sample collection stage and included in the cohort, further inspection is warranted to confirm the diagnosis. Indeed, one of the three samples (C3N-00832-T) was reclassified in the original work as pRCC upon careful examination of genomic data and was removed from the subsequent analysis.

By inspecting selected markers of RCC after removing QC samples (**Figure 2c**), we can confirm the upregulation of a typical ccRCC marker, CA9, highlighted in the original manuscript, as well as a new ccRCC marker, AHNAK2^33^. Interestingly, the novel pRCC marker PIGR^34^ is highly expressed in the outlier tumor sample (C3N-00832-T), again suggesting that this sample might be a kidney tumor of a different (non-ccRCC) subtype. A similar observation could also be made when viewing the FragPipe-Analyst generated heatmap plotted based on differentially expressed proteins (**Figure 2d**).

After excluding the outlier sample (pRCC), the results of the DE analysis based on Limma^14^ were evaluated using a volcano plot (**Figure 2e**). The comparison between tumors and NATs is similar to what previously reported in the original study^28^, with more proteins showing downregulation than upregulation in tumors versus NAT, although we applied a different statistical procedure and used only a subset of the entire sample cohort. Moreover, pathway enrichment analysis implemented in FragPipe-Analyst showed upregulation of ccRCC-related pathways, including interferon alpha response, interferon gamma response, hypoxia, mTORC1 signaling, glycolysis, and complement in tumors vs. NATs (**Figure 2d**). Oxidative phosphorylation is also downregulated, as evidenced by the low expression of NDUFV2 (**Figure2c**; **Supplementary figure S3**). These observations are in good agreement with the original study^28^.

### Analysis of a DIA-based ccRCC cancer proteomics data

Data-independent acquisition (DIA) has emerged as a widely used technology platform for quantitative protein profiling^35,36^. Several software packages have been created to support high-throughput DIA data analysis, including DIA-Umpire^37^, Skyline^38^, OpenSWATH^39^, EncyclopeDIA^40^, Spectronaut^41^, and DIA-NN^42^. We recently published our new method, MSFragger-DIA^4^. To cater to the growing interest in DIA in the field, we enabled FragPipe-Analyst to perform downstream analyses on DIA datasets processed using FragPipe (supporting both DIA-Umpire and MSFragger-DIA-based workflows). For illustration purposes, we used FragPipe-Analyst to interrogate DIA data acquired from the same ccRCC samples as described above^28^. Using quantification reports produced by DIA-NN^42^ as part of FragPipe (using the library generated by MSFragger-DIA and EasyPQP), we obtained findings highly correlated with those based on TMT data (**Supplementary figure S4, 5, 6**). Concordance was evident both at the pathway level and when comparing individual protein fold changes in tumors vs. NATs (**Supplementary figure S7**), albeit with a higher degree of ratio compression^43^ in TMT data compared to DIA. Overall, these analyses validated the results produced by FragPipe coupled to FragPipe-Analyst when applied to DIA quantification data.

### Analysis of LFQ AP-MS data

FragPipe-Analyst supports the DDA-based LFQ quantification workflows available in FragPipe, such as the LFQ-MBR workflow. For these workflows, the output quantification reports created by FragPipe/IonQuant (combined_protein.tsv) contain three quantification measures: Intensity (computed as the sum of the top N precursors for each protein), MaxLFQ intensity, and Spectral Count. The intensity (calculated using IonQuant^1^) is set as the default option in FragPipe-Analyst and is recommended for a variety of studies, including typical proteome profiling datasets, such as the ccRCC study discussed above. However, the choice of quantification method, as well as various downstream analysis options, should be assessed on a case-by-case basis, and FragPipe-Analyst provides a convenient way to perform such analyses. Here, we used FragPipe-Analyst to analyze an affinity purification mass spectrometry (AP-MS) dataset performed on a head and neck squamous cell carcinoma (HNSCC) cell line by Swaney et al.^44^. The original study aimed to reveal the changes in protein interactions due to different mutations in HNSCC. Here, we selected a subset of data from that study for illustration: the wild-type CCND1 bait in the SCC-25 cell line (three biological replicates) and four negative control runs. Importantly, for experiments where drastically different samples are compared (bait vs. negative controls), using Intensity as the quantitative measurement is preferred over MaxLFQ. In addition, imputation is important because true interactors of the bait are expected to be absent in the negative control runs. Thus, their signal is expected to be missing in the control group, and such proteins would not be captured as statistically significant without imputation. However, choosing an improper imputation method could be detrimental to the results and thus requires careful examination.

FragPipe-Analyst provides multiple imputation methods as well as visual methods to quickly assess the effects of imputation (e.g., protein abundance distributions or PCA plots, with and without imputation) and normalization (**Supplementary figure S8**). The default option is Perseus-style imputation^7^ for DDA and DIA LFQ data (no imputation is set by default for TMT data). The results of the DE analysis based on the imputed AP-MS data are illustrated by the volcano plot shown in **Figure 3a**. It shows the expected interactors, including CDK2, CDK5, CDK6, CDKN1A, CDKN1B, and CDKN1C, recapitulating the findings of the original study^44^. Furthermore, we compared the results of FragPipe-Analyst, which uses limma-based differential expression analysis, with SAINTExpress^27^, a well-established tool for AP-MS analysis previously developed by our lab (see **Supplementary figure S9**). We observed good agreement with SAINTexpress, except for ubiquitin conjugating enzyme E2 D3 (UBE2D3) and proliferating cell nuclear antigen (PCNA). Interestingly, CCND1 was previously reported to be a target of UBE2D3^45^ and to interact with PCNA^46^. As weaker or transient interactors may or may not pass statistical cutoffs depending on the specific analysis options, FragPipe-Analyst provides a convenient and complementary way for users to explore the data.

**Figure 3.**
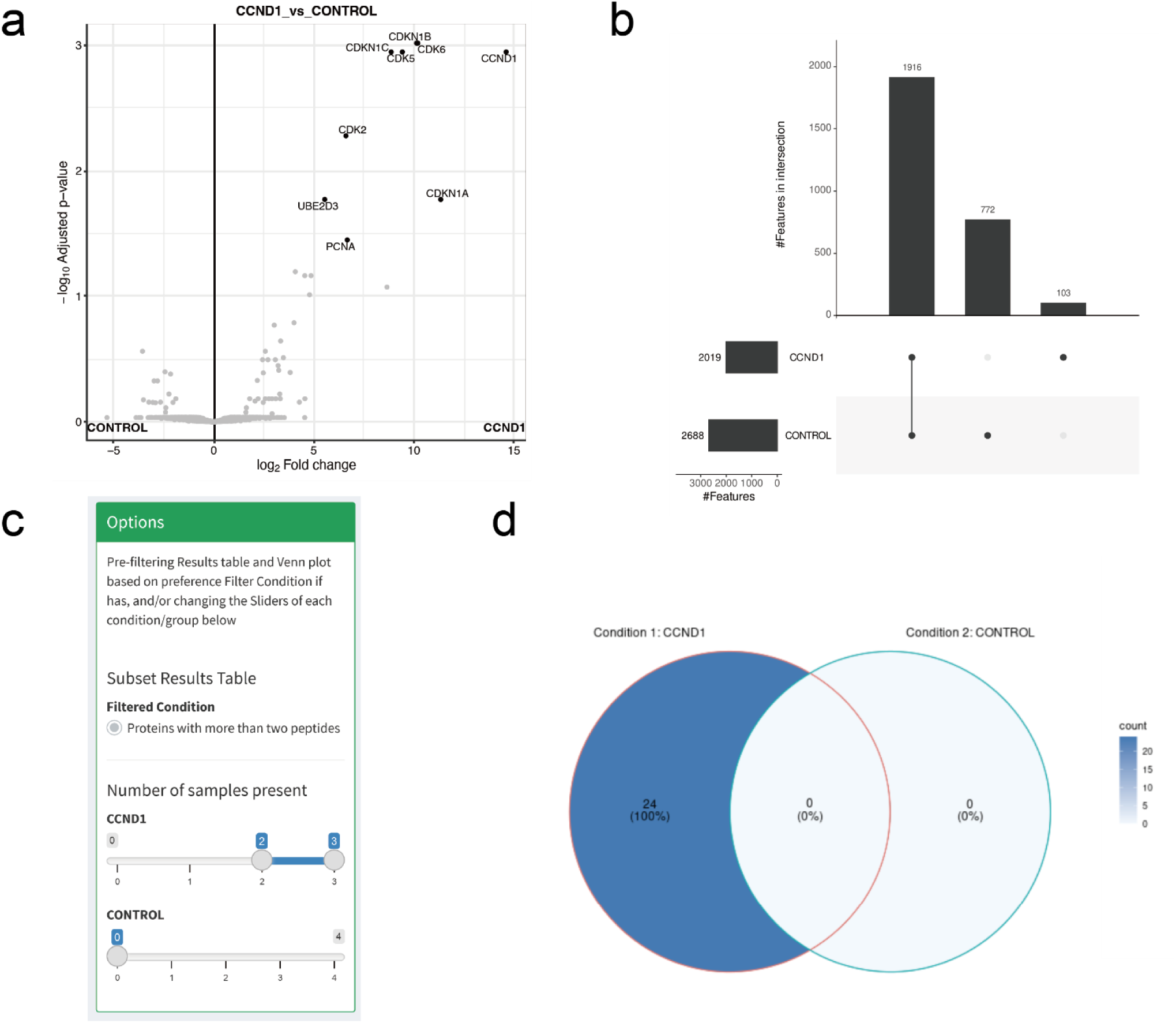
**(a)** Volcano plot comparing effect of CCND1 bait versus control. **(b)** Upset plot shows number of identified proteins across different conditions. **(c)** Screenshot of the filtering panel of the presence/absence tab and the Venn diagram **(d)** after filtering.

FragPipe-Analysis also provides users with tools to directly compare protein identifications between different conditions , without any imputation using the absence/presence tab. In the absence/presence tab, a panel with filtering options is provided to allow users to filter the protein/peptide identification table to achieve the desired confidence (**Figure 3c**). The resulting table is shown in the upper right corner of the page, and a comparison between conditions is summarized as a Venn diagram or an upset plot at the bottom (**Figure 3d**). While the Venn diagram is tailored for interpreting the overlap between two or three conditions, it may be easier to use upset plots when comparing more than three conditions.

### Peptide-level Analysis

Bottom-up proteomics usually involves quantification at the single gene/protein level by aggregating multiple peptide quantities into one to streamline interpretation and integration with other data types^13^. However, the complexity of both RNA splicing and protein post-translational modifications makes such aggregation overly simplified^13^. Several computational methods, such as PeCorA^47^, gpGrouper^48^, COPF^49^, and SEPepQuant^50^ have been recently proposed to deal with this issue, and some of them have already been built on results generated by FragPipe^47, 50^. Thus, we extended FragPipe-Analyst to support the peptide-level analysis. Aside from aggregating quantification measurements at the gene/protein level, there are several proteomics applications that require interpretation of the data at the peptide level. One such example is structural proteomics analysis using limited proteolysis coupled with mass spectrometry (LiP-MS). In a typical LiP-MS experiment, a sequence-unspecific protease, typically proteinase K (PK), is used to preferentially cleave flexible and accessible regions of the protein of interest. Since the binding of small molecules usually causes protein structural changes and can change protein activity dramatically, PK accessibility changes quantified by LiP-MS represent the readout of small molecules’ impact on the protein of interest. In addition to experimental procedures, computational tools specializing in LiP-MS have also been developed recently^51, 52^. Here, we re-analyzed a representative LiP-MS dataset of HEK293 cells treated with rapamycin^53^. The dataset consisted of 8 samples (two conditions and four biological replicates) analyzed using DIA mass spectrometry. Using the DIA_SpecLib_Quant workflow available in FragPipe, we quantified 59,503 peptides from 65,654 precursor ions across 8 samples (see **Methods**). As shown in **Figure 4**, several peptides of the known rapamycin target FKBP1A showed abundance differences in the rapamycin treatment group (RPM) compared with the DMSO control. The volcano plot of FragPipe-Analyst was designed to show not only the peptides selected in the result table (red), but also other peptides mapped to the same protein (blue). A similar peptide-level analysis can be performed using TMT data (as illustrated using the TMT ccRCC dataset; see **Supplementary figure S10, 11, and 12**). Moving forward, we plan to further develop FragPipe-Analyst to provide more advanced support for peptide-level analysis and data visualization.

**Figure 4.**
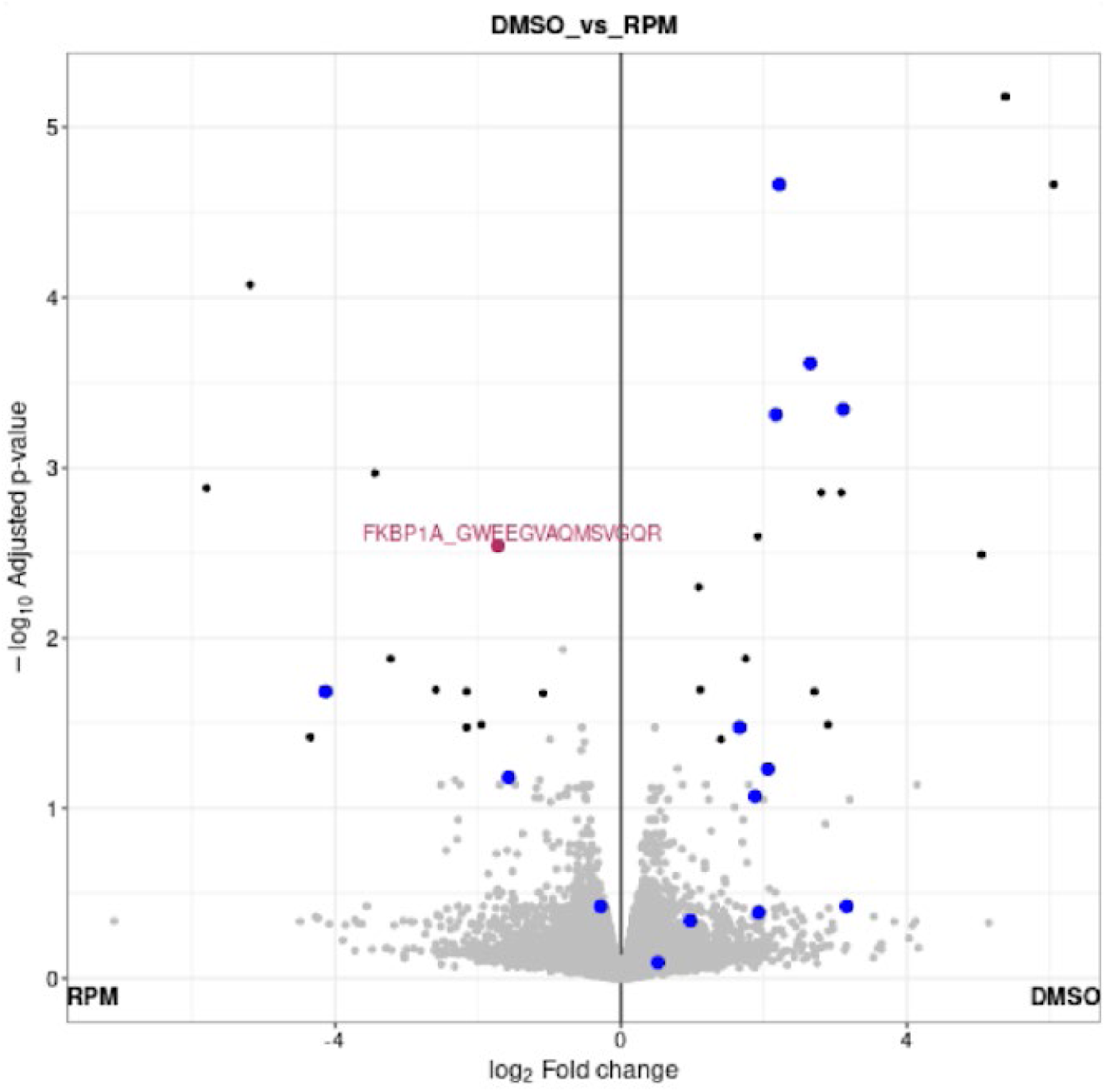
Volcano plot showing comparison of peptide intensities between rapamycin treated (RPM) and DMSO control. One peptide of FKBP1A (GWEEGVAQMSVGQR) is highlighted in red because it’s selected in the result table. Other peptides of FKBP1A are colored in blue.

### Advanced Functionality in FragPipeAnalystR: Site-Specific PTM Analysis

Large-scale proteogenomics studies, such as those by the Proteomic Tumor Analysis Consortium^28,34^ usually come with more complex experimental designs and downstream biological analysis across multi-omics data. To support more sophisticated and advanced analyses, we developed FragPipeAnalystR, an R package for FragPipe downstream analysis. It contains all functionalities available in FragPipe-Analyst but is more flexible and customizable for advanced users. Site-specific post-translational modification (PTM) analysis is a common task in proteogenomics studies. For example, protein phosphorylation is a common PTM and is important for understanding cell signaling regulation^54^. Another PTM that has received growing attention in the proteomic field is glycosylation, which plays a key role in diverse biological processes, such as cell-cell communications and immunity^55^.

Here, we illustrate FragPipeAnalystR using the TMT phosphoproteomics dataset available as part of the CPTAC ccRCC study discussed above^28^. The overall analysis strategy is illustrated in **Figure 5a**. Given that sustained proliferative signaling is a hallmark of cancers^56, 57^, it is expected that there is a large difference in protein phosphorylation between tumors and NATs, as evident from the PCA plot (**Figure 5b**). Protein phosphorylation abundance measurements are naturally correlated with whole protein abundance. To decouple changes in specific site phosphorylation levels vs. whole protein abundance changes, we built a regression model to perform normalization and take the residual protein phosphorylation to investigate biological differences between ccRCC tumors and NATs. Protein phosphorylation still differed substantially between tumors and NATs after normalization was applied (**Supplementary figure S13**). Checking specific phosphorylation sites highlighted in the original publication^28^ (PKM:P14618_Y148), MAPK1:P28482_Y187, and EIF4EBP1:Q13541_S65) confirmed their dysregulated phosphorylation in tumors (**Figure 5c**). The site-specific phosphorylation abundance differences between tumor and NAT remained after normalization for two of the three sites (MAPK1 and EIF4EBP1). These sites are of interest for studying dysregulation of cell signaling, as phosphorylation aberrations in these cases are not simply driven by changes in the abundance of the parent protein.

**Figure 5.**
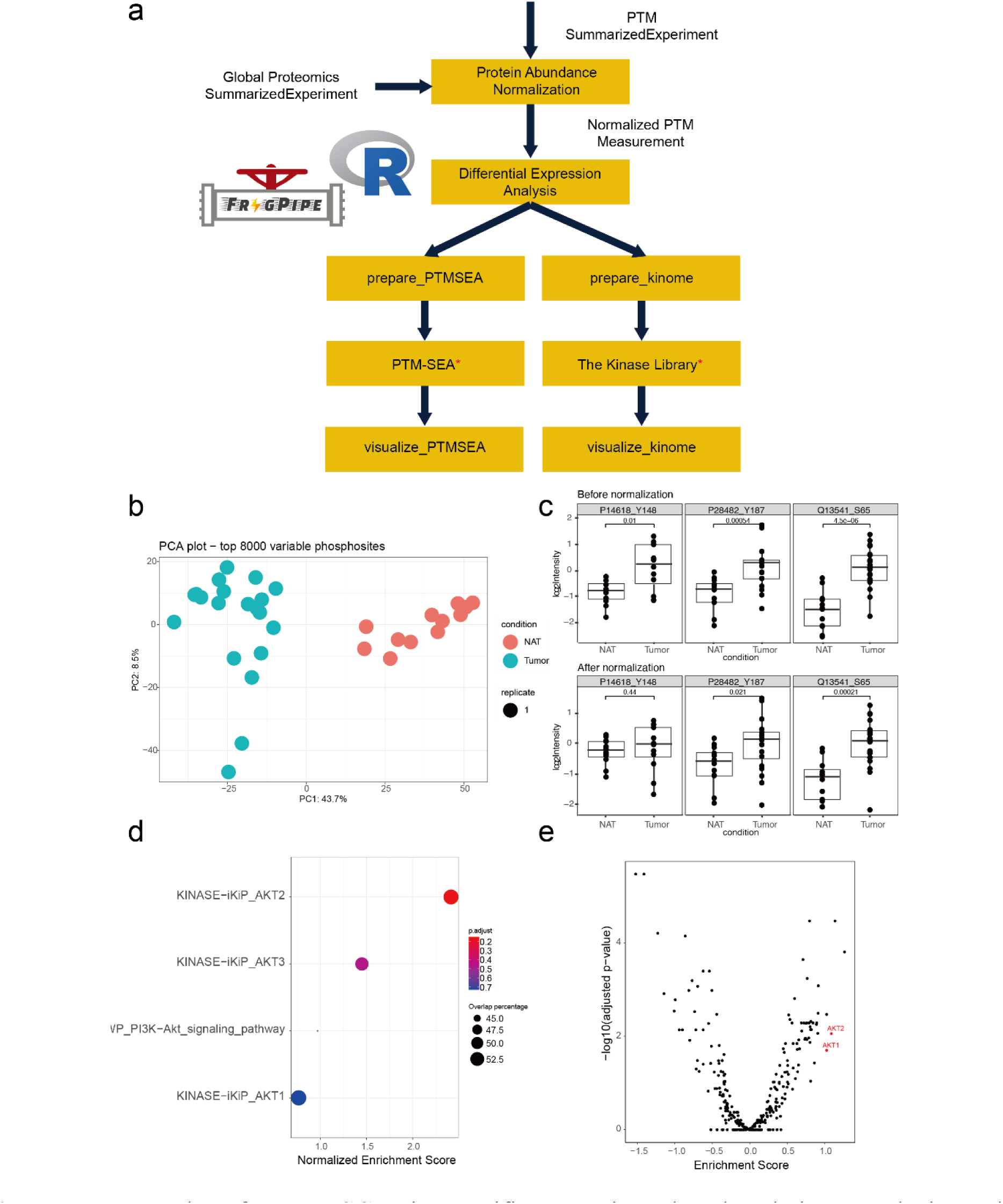
Results for ccRCC site-specific protein phosphorylation analysis using FragPipeAnalystR **(a)** The flow diagram of data processing **(b)** PCA plot of protein phosphorylation sites after normalization. **(c)** Boxplot shows abundance of selected phosphorylation sites PKM (P14618_Y148), MAPK1(P28482_Y187), and EIF4EBP1 (Q13541_S65) before and after protein abundance normalization. Results of site-specific enrichment analysis of PTM-SEA **(d)** and the Kinase library **(e)**. Both results showed the activation of AKT in ccRCC.

Site-specific PTM experiments usually generate long lists of modified sites that contain both known and novel modification sites. The interpretation of these data often becomes complicated because prior knowledge regarding the biological role of specific PTM events is limited, especially at site-level resolution. Enrichment analysis is one of the tools at our disposal for using large-scale data to generate testable hypotheses that can be investigated in a more focused manner. FragPipe-AnalystR can perform several types of enrichment analyses to support such studies. In addition to the usual enrichment analysis (i.e., using the accession number of the parent protein for each modification site), we implemented features to allow users to perform downstream site-specific PTM enrichment analyses via PTM-SEA^58^ and the Kinase library (https://kinase-library.phosphosite.org/)^59^. Results from both PTM-centric analyses showed upregulated enrichment of phosphosites associated with the AKT serine/threonine kinase family (**Figure 5d, e**). Indeed, the roles of AKT family members in RCC have already been reported^44, 60^.

## CONCLUSIONS

We presented downstream analysis software, FragPipe-Analyst, for the FragPipe user community to seamlessly perform downstream analysis after running FragPipe. FragPipe-Analyst supports all major quantification workflows (including DDA and DIA LFQ, and TMT) and offers a range of functionalities, including enhanced data exploration and peptide-level analysis. In addition to FragPipe-Analyst, we also introduced FragPipeAnalystR, an R package with all functionalities of FragPipe-Analyst at the core and supporting site-specific PTM enrichment analysis^58, 59^. We used SummarizedExperiment^61^, which is part of the R/Bioconductor ecosystem^62^, as the core data structure. This will enable us to incorporate other bioconductor bioinformatics packages in the future. Future developments will also include more complex tasks such as network analysis, proteogenomics^63^ and kinase-substrate analysis^64^. Two public server instances of FragPipe-Analyst are provided at http://fragpipe-analyst.nesvilab.org and https://fragpipe-analyst.org. FragPipe-Analyst and FragPipeAnalystR are open-source and freely available at https://github.com/MonashProteomics/FragPipe-Analyst and https://github.com/Nesvilab/FragPipeAnalystR, respectively. A Docker file is also included in the FragPipe-Analyst GitHub repository to help users set up their own instances without requiring the transfer of data through the Internet.

## ASSOCIATED CONTENT

### Supporting Information

The following files are available free of charge.

- Supporting information document (PDF) contains supplementary figures.
- Quantification report and experimental annotation of the ccRCC TMT global proteomics dataset
- Example usage of FragPipeAnalystR package (R script, R)

**Supplementary figure S1**. PCA plot by batch of ccRCC proteomics dataset collected by TMT.

**Supplementary figure S2**. Missing value heatmap of ccRCC proteomics dataset collected by TMT.

**Supplementary figure S3**. Over representation test result for downregulation part in tumors vs NAT comparison of ccRCC proteomics dataset collected by TMT.

**Supplementary figure S4**. PCA plot of ccRCC proteomics dataset collected by data-independent acquisition (DIA).

**Supplementary figure S5**. Correlation heatmap of ccRCC proteomics dataset collected by DIA.

**Supplementary figure S6**. Volcano plot (left) and overrepresentation test result (right) of ccRCC proteomics dataset collected by DIA.

**Supplementary figure S7**. Scatter plot shows comparison of log2 fold changes obtained from DIA and TMT experiments when comparing tumors versus NATs.

**Supplementary figure S8**. Density plot shows the imputation effect on AP-MS experiments used here.

**Supplementary figure S9**. Network visualization of HNSCC AP-MS dataset generated by SAINTexpress.

**Supplementary figure S10**. PCA plot of peptide-level analysis on ccRCC proteomics dataset.

**Supplementary figure S11**. Correlation matrix plot of ccRCC proteomics dataset using peptide-level quantification result.

**Supplementary figure S12**. Volcano plot (left) and overrepresentation test result (right) of peptide-level analysis on ccRCC proteomics dataset.

**Supplementary figure S13**. PCA plot of ccRCC phosphoproteomics dataset after normalization.

## Author Contributions

Y.H. programmed and created FragPipe-Analyst and FragPipeAnalystR, based on the initial LFQ-Analyst code. H.Z. helped with the FragPipe-Analyst feature development. G.X.L. provided suggestions for statistical procedures and helped develop FragPipe-Analyst and FragPipeAnalyst.

F.Y. provided suggestions on the input FragPipe data format and streamlined the support of annotation files in FragPipe. Y.H. drafted the first version of this manuscript. H.Z., G.X.L., J.R.S. and A. I. N. reviewed and edited the manuscript. A.I.N. and R.B.S. conceived of the study. The manuscript was written through the contributions of all authors. All the authors approved the final version of the manuscript.

## ACKNOWLEDGMENTS

This work was supported by NIH grants R01-GM-094231, U24-CA210967, and U24-CA271037 received by Y. H., G.X.L., F. Y., and A.I.N., and the Monash Proteomics & Metabolomics Platform with support from Bioplatforms Australia (BPA) as part of the National Collaborative Research Infrastructure Strategy (NCRIS).

## ABBREVIATIONS

PTM: post-translational modification
DDA: data-dependent acquisition
DIA: data-independent acquisition
LFQ: label-free quantification
RCC: renal cell carcinoma
ccRCC: clear cell renal cell carcinoma
ChRCC: chromophobe renal cell carcinoma

## REFERENCES

1. Yu, F.; Haynes, S. E.; Nesvizhskii, A. I., IonQuant enables accurate and sensitive label-free quantification with FDR-controlled match-between-runs. Molecular & Cellular Proteomics 2021, 20.

2. Djomehri, S. I.; Gonzalez, M. E.; da Veiga Leprevost, F.; Tekula, S. R.; Chang, H.-Y.; White, M. J.; Cimino-Mathews, A.; Burman, B.; Basrur, V.; Argani, P., Quantitative proteomic landscape of metaplastic breast carcinoma pathological subtypes and their relationship to triple-negative tumors. Nature communications 2020, 11 (1), 1723.

3. Demichev, V.; Szyrwiel, L.; Yu, F.; Teo, G. C.; Rosenberger, G.; Niewienda, A.; Ludwig, D.; Decker, J.; Kaspar-Schoenefeld, S.; Lilley, K. S., dia-PASEF data analysis using FragPipe and DIA-NN for deep proteomics of low sample amounts. Nature communications 2022, 13 (1), 3944.

4. Yu, F.; Teo, G. C.; Kong, A. T.; Fröhlich, K.; Li, G. X.; Demichev, V.; Nesvizhskii, A. I., Analysis of DIA proteomics data using MSFragger-DIA and FragPipe computational platform. Nature Communications 2023, 14 (1), 4154.

4. Zhang, X.; Smits, A. H.; Van Tilburg, G. B. A.; Ovaa, H.; Huber, W.; Vermeulen, M., Proteome-wide identification of ubiquitin interactions using UbIA-MS. Nature protocols 2018, 13 (3), 530–550.

5. Quast, J.-P.; Schuster, D.; Picotti, P., protti: an R package for comprehensive data analysis of peptide-and protein-centric bottom-up proteomics data. Bioinformatics Advances 2021, 2 (1).

6. Tyanova, S.; Temu, T.; Sinitcyn, P.; Carlson, A.; Hein, M. Y.; Geiger, T.; Mann, M.; Cox, J., The Perseus computational platform for comprehensive analysis of (prote)omics data. Nature Methods 2016, 13 (9), 731–740.

7. Shah, A. D.; Goode, R. J. A.; Huang, C.; Powell, D. R.; Schittenhelm, R. B., LFQ-analyst: an easy-to-use interactive web platform to analyze and visualize label-free proteomics data preprocessed with MaxQuant. Journal of proteome research 2019, 19 (1), 204–211.

8. Choi, M.; Chang, C.-Y.; Clough, T.; Broudy, D.; Killeen, T.; MacLean, B.; Vitek, O., MSstats: an R package for statistical analysis of quantitative mass spectrometry-based proteomic experiments. Bioinformatics 2014, 30 (17), 2524–2526.

9. Huang, T.; Choi, M.; Tzouros, M.; Golling, S.; Pandya, N. J.; Banfai, B.; Dunkley, T.; Vitek, O., MSstatsTMT: statistical detection of differentially abundant proteins in experiments with isobaric labeling and multiple mixtures. Molecular & Cellular Proteomics 2020, 19 (10), 1706–1723.

10. Tyanova, S.; Temu, T.; Cox, J., The MaxQuant computational platform for mass spectrometry-based shotgun proteomics. Nature protocols 2016, 11 (12), 2301–2319.

11. Bai, M.; Deng, J.; Dai, C.; Pfeuffer, J.; Sachsenberg, T.; Perez-Riverol, Y., LFQ-Based Peptide and Protein Intensity Differential Expression Analysis. Journal of Proteome Research 2023, 22 (6), 2114–2123.

12. Plubell, D. L.; Käll, L.; Webb-Robertson, B.-J.; Bramer, L. M.; Ives, A.; Kelleher, N. L.; Smith, L. M.; Montine, T. J.; Wu, C. C.; MacCoss, M. J., Putting humpty dumpty back together again: what does protein quantification mean in bottom-up proteomics? Journal of proteome research 2022, 21 (4), 891–898.

13. Smyth, G. K., Limma: linear models for microarray data. Bioinformatics and computational biology solutions using R and Bioconductor 2005, 397–420.

14. Kuleshov, M. V.; Jones, M. R.; Rouillard, A. D.; Fernandez, N. F.; Duan, Q.; Wang, Z.; Koplev, S.; Jenkins, S. L.; Jagodnik, K. M.; Lachmann, A., Enrichr: a comprehensive gene set enrichment analysis web server 2016 update. Nucleic acids research 2016, 44 (W1), W90–W97.

15. Huber, W.; Von Heydebreck, A.; Sültmann, H.; Poustka, A.; Vingron, M., Variance stabilization applied to microarray data calibration and to the quantification of differential expression. Bioinformatics 2002, 18 (suppl_1), S96–S104.

16. Gatto, L.; Lilley, K. S., MSnbase-an R/Bioconductor package for isobaric tagged mass spectrometry data visualization, processing and quantitation. Bioinformatics 2012, 28 (2), 288–289.

17. Yu, G.; Wang, L.-G.; Han, Y.; He, Q.-Y., clusterProfiler: an R package for comparing biological themes among gene clusters. Omics: a journal of integrative biology 2012, 16 (5), 284–287.

18. Wickham, H., ggplot2. Wiley interdisciplinary reviews: computational statistics 2011, 3 (2), 180–185.

19. Gu, Z.; Eils, R.; Schlesner, M., Complex heatmaps reveal patterns and correlations in multidimensional genomic data. Bioinformatics 2016, 32 (18), 2847–2849.

20. Sievert, C., Interactive web-based data visualization with R, plotly, and shiny. CRC Press: 2020.

21. Kong, A. T.; Leprevost, F. V.; Avtonomov, D. M.; Mellacheruvu, D.; Nesvizhskii, A. I., MSFragger: ultrafast and comprehensive peptide identification in mass spectrometry–based proteomics. Nature methods 2017, 14 (5), 513–520.

22. Yu, F.; Teo, G. C.; Kong, A. T.; Haynes, S. E.; Avtonomov, D. M.; Geiszler, D. J.; Nesvizhskii, A. I., Identification of modified peptides using localization-aware open search. Nature communications 2020, 11 (1), 4065.

23. Yang, K. L.; Yu, F.; Teo, G. C.; Li, K.; Demichev, V.; Ralser, M.; Nesvizhskii, A. I., MSBooster: improving peptide identification rates using deep learning-based features. Nature Communications 2023, 14 (1), 4539.

24. Käll, L.; Canterbury, J. D.; Weston, J.; Noble, W. S.; MacCoss, M. J., Semi-supervised learning for peptide identification from shotgun proteomics datasets. Nature methods 2007, 4 (11), 923–925.

25. da Veiga Leprevost, F.; Haynes, S. E.; Avtonomov, D. M.; Chang, H.-Y.; Shanmugam, A. K.; Mellacheruvu, D.; Kong, A. T.; Nesvizhskii, A. I., Philosopher: a versatile toolkit for shotgun proteomics data analysis. Nature methods 2020, 17 (9), 869–870.

26. Teo, G.; Liu, G.; Zhang, J.; Nesvizhskii, A. I.; Gingras, A.-C.; Choi, H., SAINTexpress: improvements and additional features in Significance Analysis of INTeractome software. Journal of proteomics 2014, 100, 37–43.

27. Clark, D. J.; Dhanasekaran, S. M.; Petralia, F.; Pan, J.; Song, X.; Hu, Y.; da Veiga Leprevost, F.; Reva, B.; Lih, T.-S. M.; Chang, H.-Y., Integrated proteogenomic characterization of clear cell renal cell carcinoma. Cell 2019, 179 (4), 964–983. e31.

28. Consortium, U., UniProt: a worldwide hub of protein knowledge. Nucleic acids research 2019, 47 (D1), D506–D515.

29. Shteynberg, D. D.; Deutsch, E. W.; Campbell, D. S.; Hoopmann, M. R.; Kusebauch, U.; Lee, D.; Mendoza, L.; Midha, M. K.; Sun, Z.; Whetton, A. D., PTMProphet: Fast and accurate mass modification localization for the trans-proteomic pipeline. Journal of proteome research 2019, 18 (12), 4262–4272.

30. Nesvizhskii, A. I.; Keller, A.; Kolker, E.; Aebersold, R., A statistical model for identifying proteins by tandem mass spectrometry. Analytical chemistry 2003, 75 (17), 4646–4658.

31. Johnson, W. E.; Li, C.; Rabinovic, A., Adjusting batch effects in microarray expression data using empirical Bayes methods. Biostatistics 2007, 8 (1), 118–127.

32. Wang, M.; Li, X.; Zhang, J.; Yang, Q.; Chen, W.; Jin, W.; Huang, Y.-R.; Yang, R.; Gao, W.-Q., AHNAK2 is a novel prognostic marker and oncogenic protein for clear cell renal cell carcinoma. Theranostics 2017, 7 (5), 1100.

33. Li, G. X.; Hsiao, Y.; Chen, L.; Mannan, R.; Zhang, Y.; Petralia, F.; Cho, H.; Hosseini, N.; Calinawan, A.; Li, Y.; Anand, S.; Dagar, A.; Geffen, Y.; Leprevost, F. V.; Le, A.; Ponce, S.; Schnaubelt, M.; Deen, N. N. A.; Caravan, W.; Houston, A.; Kumar-Sinha, C.; Wang, X.; Chugh, S.; Omenn, G. S.; Chan, D. W.; Ricketts, C.; Mehra, R.; Chinnaiyan, A.; Ding, L.; Cieslik, M.; Zhang, H.; Dhanasekaran, S. M.; Nesvizhskii, A. I., Abstract 3127: Comprehensive proteogenomic characterization of rare kidney tumors. Cancer Research 2023, 83 (7_Supplement), 3127–3127.

34. Kitata, R. B.; Yang, J. C.; Chen, Y. J., Advances in data-independent acquisition mass spectrometry towards comprehensive digital proteome landscape. Mass spectrometry reviews 2023, 42 (6), 2324–2348.

35. Ludwig, C.; Gillet, L.; Rosenberger, G.; Amon, S.; Collins, B. C.; Aebersold, R., Data-independent acquisition-based SWATH-MS for quantitative proteomics: a tutorial. Molecular systems biology 2018, 14 (8), e8126.

36. Tsou, C.-C.; Avtonomov, D.; Larsen, B.; Tucholska, M.; Choi, H.; Gingras, A.-C.; Nesvizhskii, A. I., DIA-Umpire: comprehensive computational framework for data-independent acquisition proteomics. Nature methods 2015, 12 (3), 258–264.

37. MacLean, B.; Tomazela, D. M.; Shulman, N.; Chambers, M.; Finney, G. L.; Frewen, B.; Kern, R.; Tabb, D. L.; Liebler, D. C.; MacCoss, M. J., Skyline: an open source document editor for creating and analyzing targeted proteomics experiments. Bioinformatics 2010, 26 (7), 966–968.

38. Röst, H. L.; Rosenberger, G.; Navarro, P.; Gillet, L.; Miladinović, S. M.; Schubert, O. T.; Wolski, W.; Collins, B. C.; Malmström, J.; Malmström, L., OpenSWATH enables automated, targeted analysis of data-independent acquisition MS data. Nature biotechnology 2014, 32 (3), 219–223.

39. Searle, B. C.; Swearingen, K. E.; Barnes, C. A.; Schmidt, T.; Gessulat, S.; Küster, B.; Wilhelm, M., Generating high quality libraries for DIA MS with empirically corrected peptide predictions. Nature communications 2020, 11 (1), 1548.

40. Bruderer, R.; Bernhardt, O. M.; Gandhi, T.; Miladinović, S. M.; Cheng, L.-Y.; Messner, S.; Ehrenberger, T.; Zanotelli, V.; Butscheid, Y.; Escher, C., Extending the limits of quantitative proteome profiling with data-independent acquisition and application to acetaminophen-treated three-dimensional liver microtissues*[S]. Molecular & Cellular Proteomics 2015, 14 (5), 1400–1410.

41. Demichev, V.; Messner, C. B.; Vernardis, S. I.; Lilley, K. S.; Ralser, M., DIA-NN: neural networks and interference correction enable deep proteome coverage in high throughput. Nature methods 2020, 17 (1), 41–44.

42. Savitski, M. M.; Mathieson, T.; Zinn, N.; Sweetman, G.; Doce, C.; Becher, I.; Pachl, F.; Kuster, B.; Bantscheff, M., Measuring and managing ratio compression for accurate iTRAQ/TMT quantification. Journal of proteome research 2013, 12 (8), 3586–3598.

43. Swaney, D. L.; Ramms, D. J.; Wang, Z.; Park, J.; Goto, Y.; Soucheray, M.; Bhola, N.; Kim, K.; Zheng, F.; Zeng, Y., A protein network map of head and neck cancer reveals PIK3CA mutant drug sensitivity. Science 2021, 374 (6563), eabf2911.

44. Hattori, H.; Zhang, X.; Jia, Y.; Subramanian, K. K.; Jo, H.; Loison, F.; Newburger, P. E.; Luo, H. R., RNAi screen identifies UBE2D3 as a mediator of all-trans retinoic acid-induced cell growth arrest in human acute promyelocytic NB4 cells. Blood 2007, 110 (2), 640–650.

45. Matsuoka, S.; Yamaguchi, M.; Matsukage, A., D-type cyclin-binding regions of proliferating cell nuclear antigen. Journal of Biological Chemistry 1994, 269 (15), 11030–11036.

46. Dermit, M.; Peters-Clarke, T. M.; Shishkova, E.; Meyer, J. G., Peptide correlation analysis (PeCorA) reveals differential proteoform regulation. Journal of proteome research 2020, 20 (4), 1972–1980.

47. Saltzman, A. B.; Leng, M.; Bhatt, B.; Singh, P.; Chan, D. W.; Dobrolecki, L.; Chandrasekaran, H.; Choi, J. M.; Jain, A.; Jung, S. Y., gpGrouper: a peptide grouping algorithm for gene-centric inference and quantitation of bottom-up proteomics data. Molecular & Cellular Proteomics 2018, 17 (11), 2270–2283.

48. Bludau, I.; Frank, M.; Dörig, C.; Cai, Y.; Heusel, M.; Rosenberger, G.; Picotti, P.; Collins, B. C.; Röst, H.; Aebersold, R., Systematic detection of functional proteoform groups from bottom-up proteomic datasets. Nature communications 2021, 12 (1), 3810.

49. Dou, Y.; Liu, Y.; Yi, X.; Olsen, L. K.; Zhu, H.; Gao, Q.; Zhou, H.; Zhang, B., SEPepQuant enhances the detection of possible isoform regulations in shotgun proteomics. Nature Communications 2023, 14 (1), 5809.

50. Manriquez-Sandoval, E.; Brewer, J.; Lule, G.; Lopez, S.; Fried, S. D., FLiPPR: A Processor for Limited Proteolysis (LiP) Mass Spectrometry Datasets Built on FragPipe. bioRxiv 2023, 2023.12.04.569947.

51. Piazza, I.; Beaton, N.; Bruderer, R.; Knobloch, T.; Barbisan, C.; Chandat, L.; Sudau, A.; Siepe, I.; Rinner, O.; de Souza, N., A machine learning-based chemoproteomic approach to identify drug targets and binding sites in complex proteomes. Nature Communications 2020, 11 (1), 4200.

52. Reber, V.; Gstaiger, M., Target Deconvolution by Limited Proteolysis Coupled to Mass Spectrometry. In Chemogenomics, Merk, D.; Chaikuad, A., Eds. Springer US: New York, NY, 2023; Vol. 2706, pp 177–190.

53. Pawson, T.; Scott, J. D., Protein phosphorylation in signaling – 50 years and counting. Trends in Biochemical Sciences 2005, 30 (6), 286–290.

54. Bagdonaite, I.; Malaker, S. A.; Polasky, D. A.; Riley, N. M.; Schjoldager, K.; Vakhrushev, S. Y.; Halim, A.; Aoki-Kinoshita, K. F.; Nesvizhskii, A. I.; Bertozzi, C. R., Glycoproteomics. Nature Reviews Methods Primers 2022, 2 (1), 48.

55. Hanahan, D.; Weinberg, R. A., The hallmarks of cancer. cell 2000, 100 (1), 57–70.

56. Hanahan, D.; Weinberg, R. A., Hallmarks of cancer: the next generation. cell 2011, 144 (5), 646–674.

57. Krug, K.; Mertins, P.; Zhang, B.; Hornbeck, P.; Raju, R.; Ahmad, R.; Szucs, M.; Mundt, F.; Forestier, D.; Jane-Valbuena, J., A Curated Resource for Phosphosite-specific Signature Analysis*[S]. Molecular & cellular proteomics 2019, 18 (3), 576–593.

58. Johnson, J. L.; Yaron, T. M.; Huntsman, E. M.; Kerelsky, A.; Song, J.; Regev, A.; Lin, T.-Y.; Liberatore, K.; Cizin, D. M.; Cohen, B. M., An atlas of substrate specificities for the human serine/threonine kinome. Nature 2023, 1–8.

59. Guo, H.; German, P.; Bai, S.; Barnes, S.; Guo, W.; Qi, X.; Lou, H.; Liang, J.; Jonasch, E.; Mills, G. B., The PI3K/AKT pathway and renal cell carcinoma. Journal of genetics and genomics 2015, 42 (7), 343–353.

60. Martin Morgan, V. O. SummarizedExperiment, Bioconductor: 2017.

61. Gentleman, R. C.; Carey, V. J.; Bates, D. M.; Bolstad, B.; Dettling, M.; Dudoit, S.; Ellis, B.; Gautier, L.; Ge, Y.; Gentry, J., Bioconductor: open software development for computational biology and bioinformatics. Genome biology 2004, 5 (10), 1–16.

62. Nesvizhskii, A. I., Proteogenomics: concepts, applications and computational strategies. Nature Methods 2014, 11 (11), 1114–1125.

63. Han, B.; Li, G. X.; Liew, W. L.; Chan, E.; Huang, S.; Khoo, C. M.; Leow, M. K.-S.; Lee, Y. S.; Zhao, T.; Wang, L. C.; Sobota, R.; Choi, H.; Liu, M. H.; Kim, K. P.; Tai, E. S., Unbiased phosphoproteomics analysis unveils modulation of insulin signaling by extramitotic CDK1 kinase activity in human myotubes. bioRxiv 2023, 2023.06.30.547176.

