## Supplementary Information for "Analysis and visualization of quantitative proteomics data using FragPipe-Analyst"

**Supplementary figure S1**. PCA plot by batch of ccRCC proteomics dataset collected by TMT.
**Supplementary figure S2**. Missing value heatmap of ccRCC proteomics dataset collected by TMT.
**Supplementary figure S3**. Over representation test result for downregulation part in tumors vs NAT comparison of ccRCC proteomics dataset collected by TMT.
**Supplementary figure S4**. PCA plot of ccRCC proteomics dataset collected by data-independent acquisition (DIA).
**Supplementary figure S5**. Correlation heatmap of ccRCC proteomics dataset collected by DIA.
**Supplementary figure S6**. Volcano plot (left) and overrepresentation test result (right) of ccRCC proteomics dataset collected by DIA.

**Supplementary figure S7**. Scatter plot shows comparison of log2 fold changes obtained from DIA and TMT experiments when comparing tumors versus NATs.
**Supplementary figure S8**. Density plot shows the imputation effect on AP-MS experiments used here.
**Supplementary figure S9**. Network visualization of HNSCC AP-MS dataset generated by SAINTexpress.
**Supplementary figure S10.** PCA plot of peptide-level analysis on ccRCC proteomics dataset.
**Supplementary figure S11**. Correlation matrix plot of ccRCC proteomics dataset using peptide-level quantification result.
**Supplementary figure S12**. Volcano plot (left) and overrepresentation test result (right) of peptide-level analysis on ccRCC proteomics dataset.
**Supplementary figure S13**. PCA plot of ccRCC phosphoproteomics dataset after normalization.

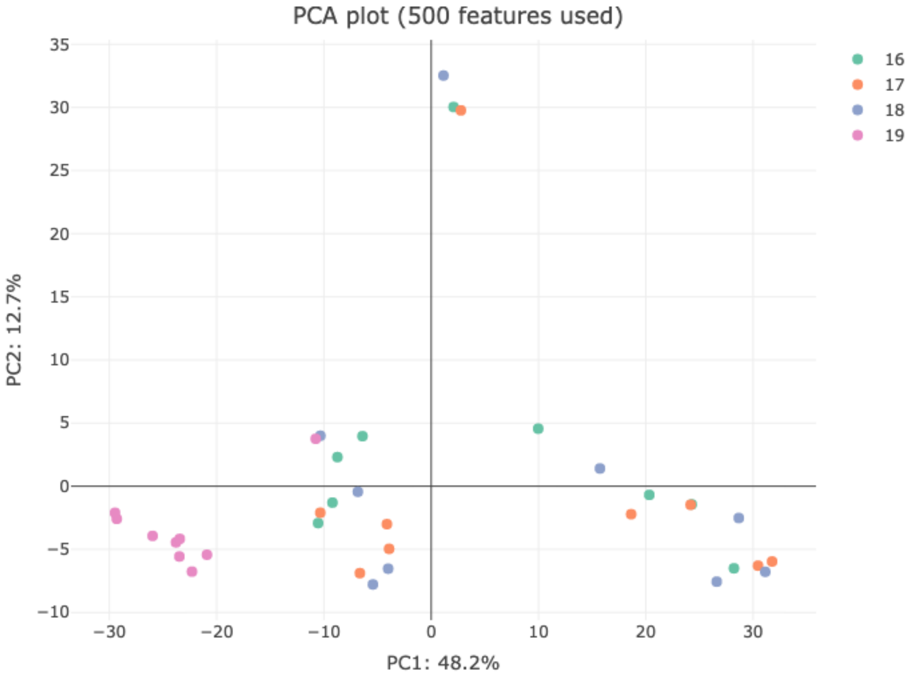


**Supplementary figure S1**. PCA plot by batch of ccRCC proteomics dataset collected by TMT.

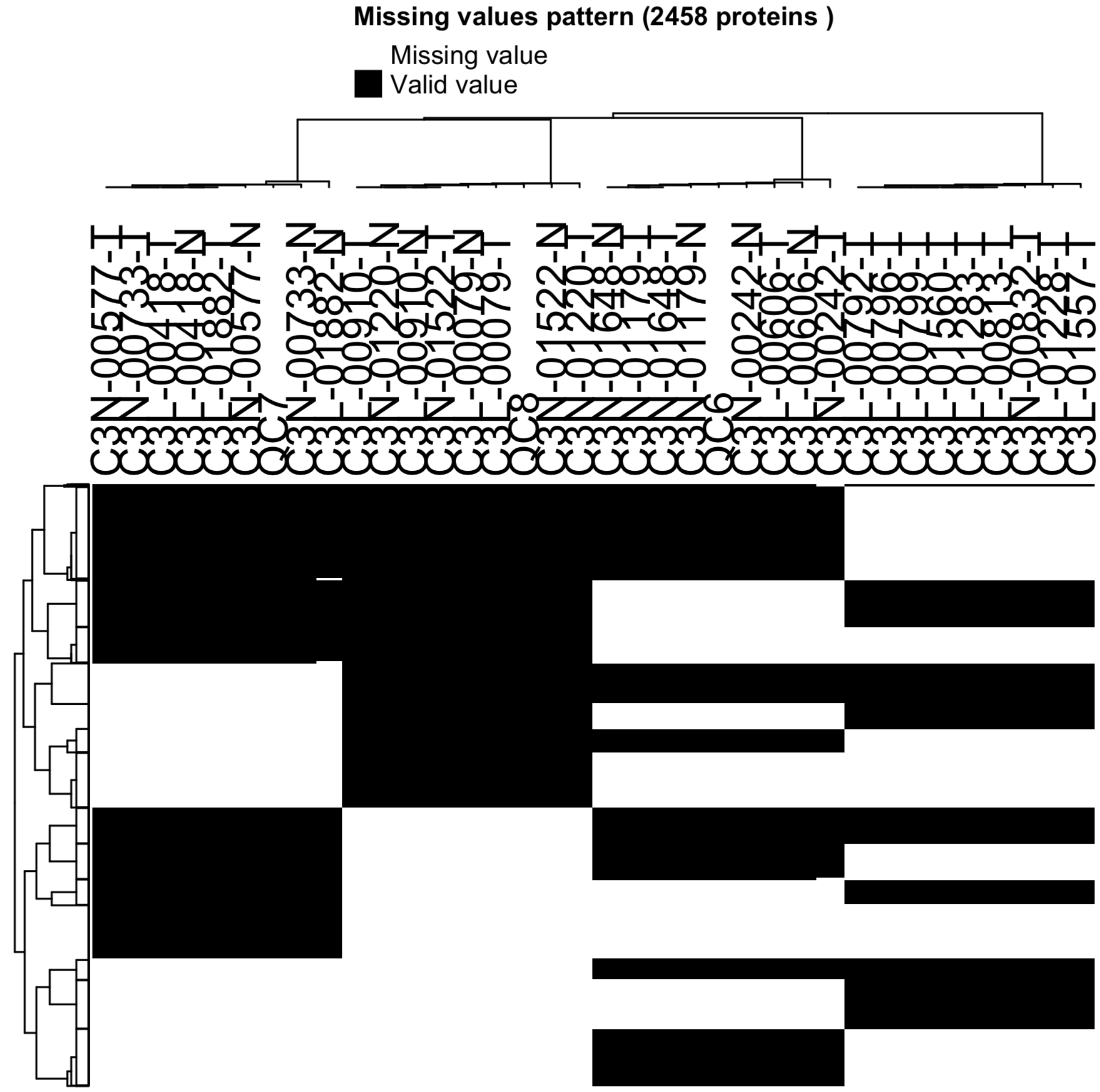


**Supplementary figure S2**. Missing value heatmap of ccRCC proteomics dataset collected by TMT.

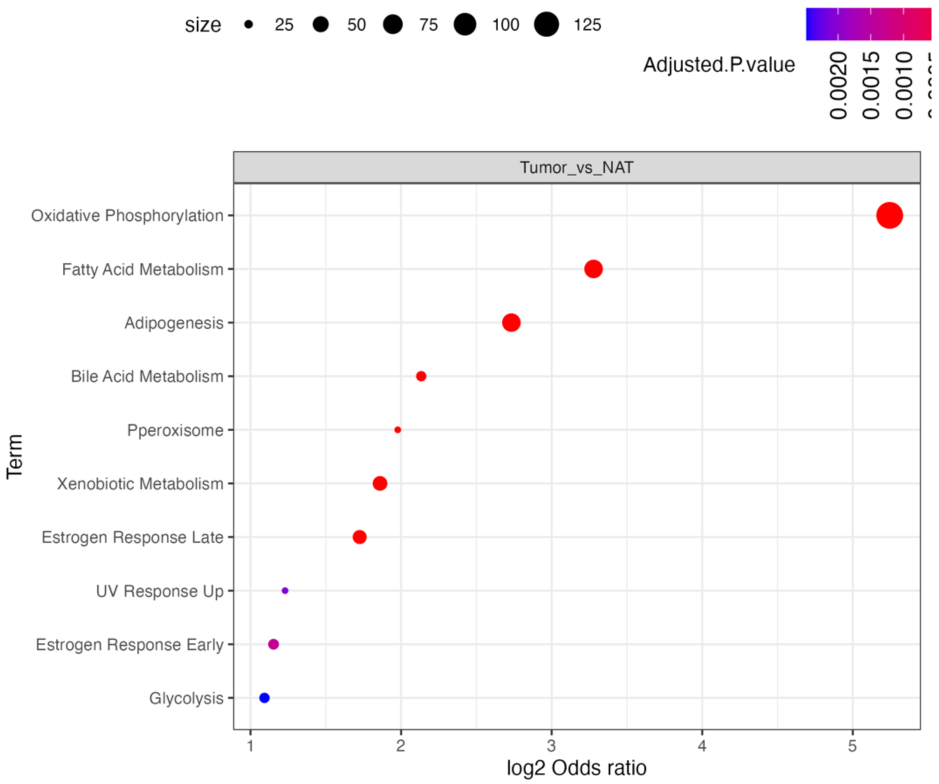


**Supplementary figure S3**. Over representation test result for downregulation part in tumors vs NAT comparison of ccRCC proteomics dataset collected by TMT.

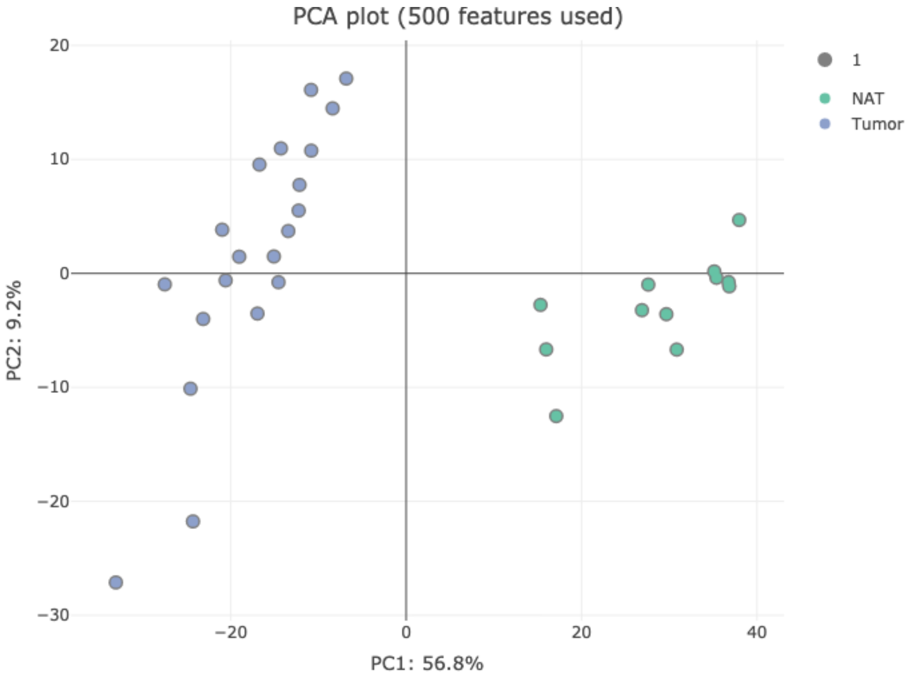


**Supplementary figure S4**. PCA plot of ccRCC proteomics dataset collected by data-independent acquisition (DIA).

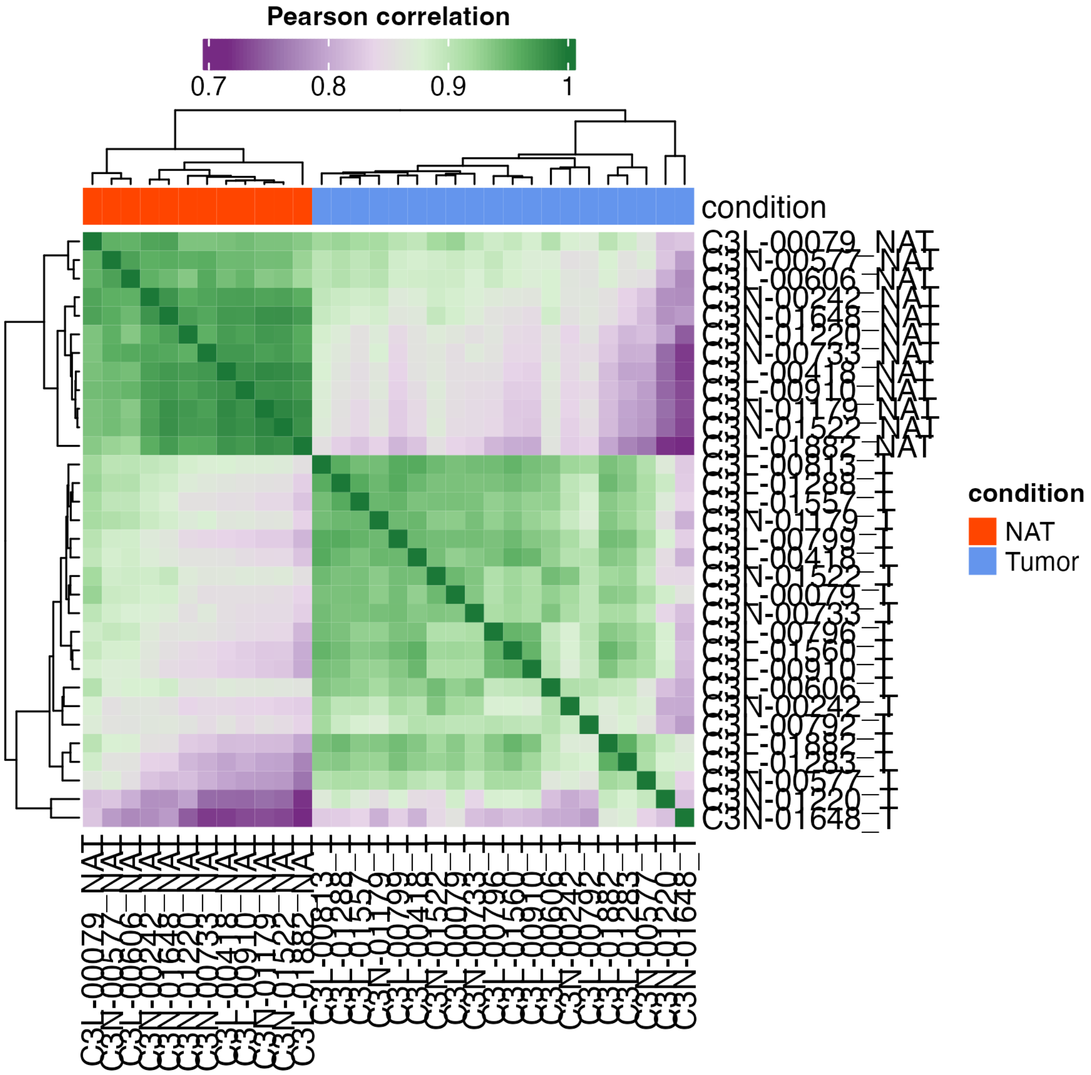


**Supplementary figure S5**. Correlation heatmap of ccRCC proteomics dataset collected by DIA.

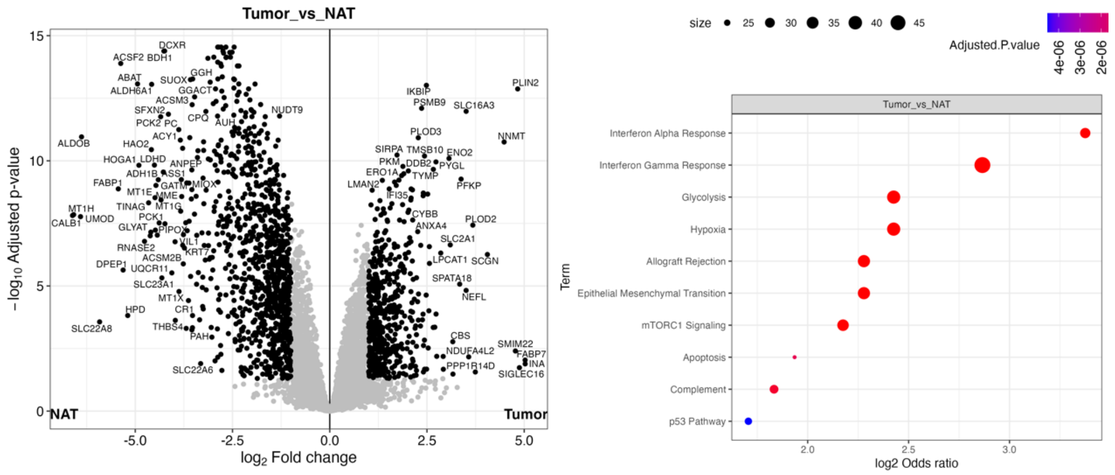


**Supplementary figure S6**. Volcano plot (left) and overrepresentation test result (right) of ccRCC proteomics dataset collected by DIA.

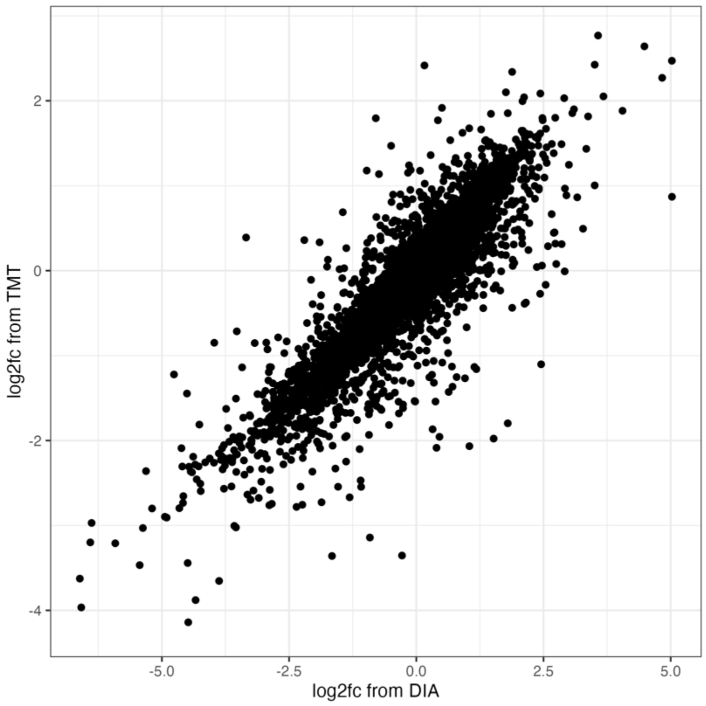


**Supplementary figure S7**. Scatter plot shows comparison of log2 fold changes obtained from DIA and TMT experiments when comparing tumors versus NATs.

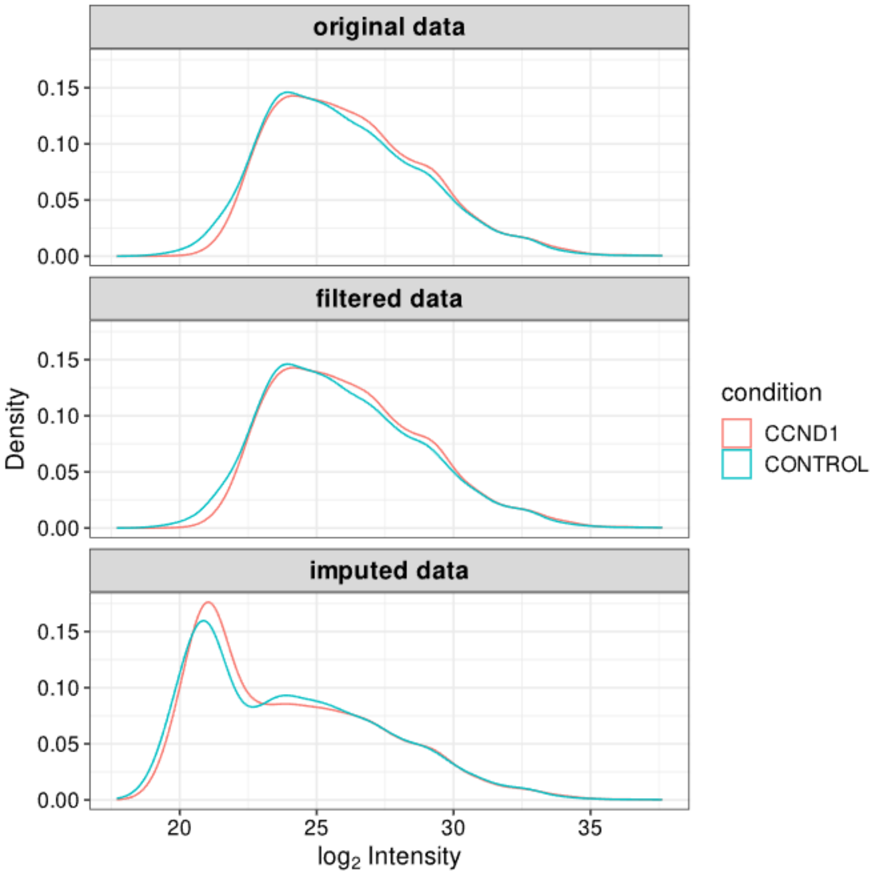


**Supplementary figure S8**. Density plot shows the imputation effect (Perseus style imputation) on AP-MS experiments.

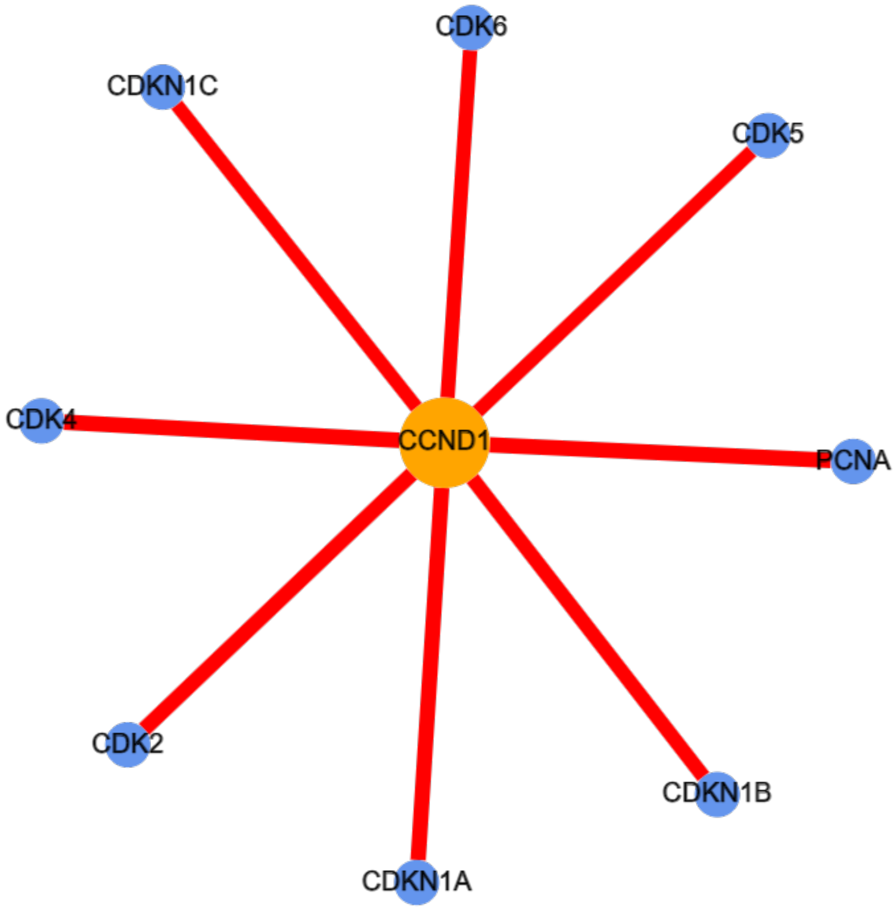


**Supplementary figure S9**. Network visualization of HNSCC AP-MS dataset generated by SAINTexpress.

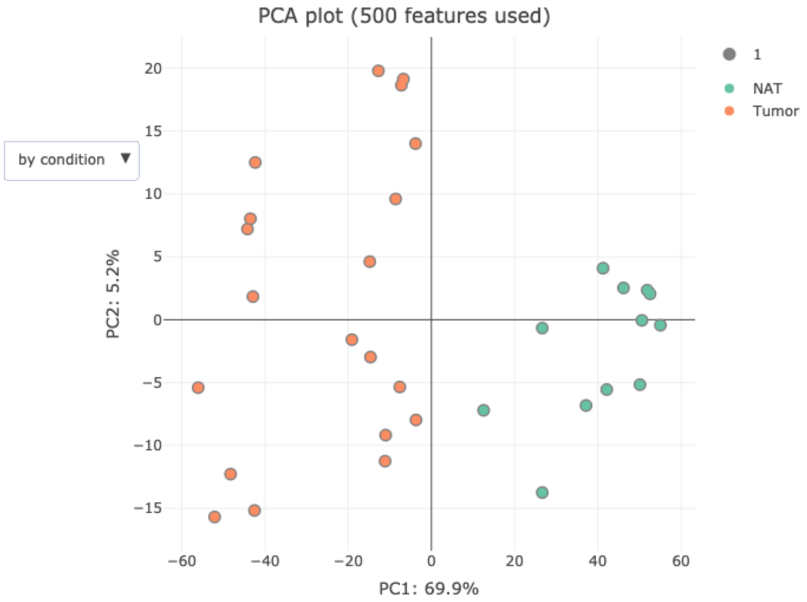


**Supplementary figure S10**. PCA plot of peptide-level analysis of ccRCC proteomics dataset.

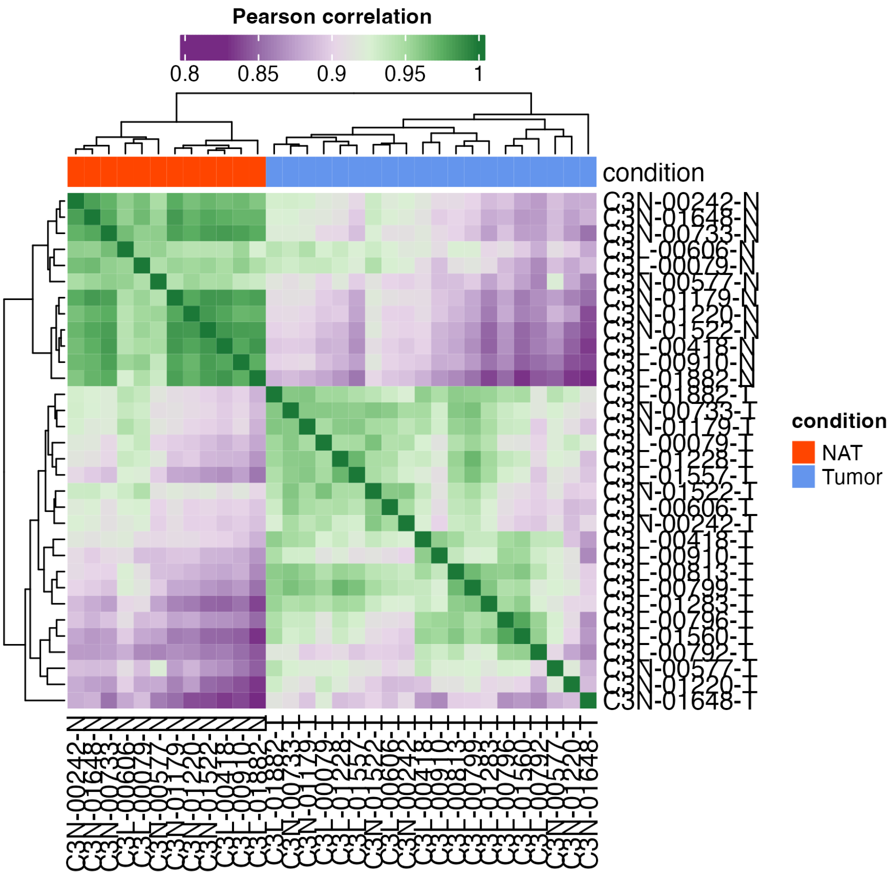


**Supplementary figure S11**. Correlation matrix plot of ccRCC proteomics dataset using peptide-level quantification result.

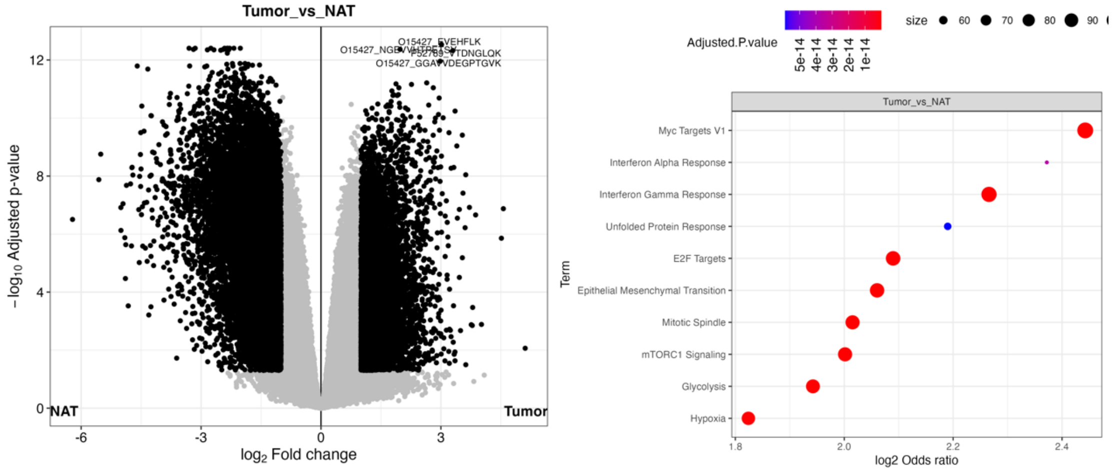


**Supplementary figure S12**. Volcano plot (left) and overrepresentation test result (right) of peptide-level analysis on ccRCC proteomics dataset.

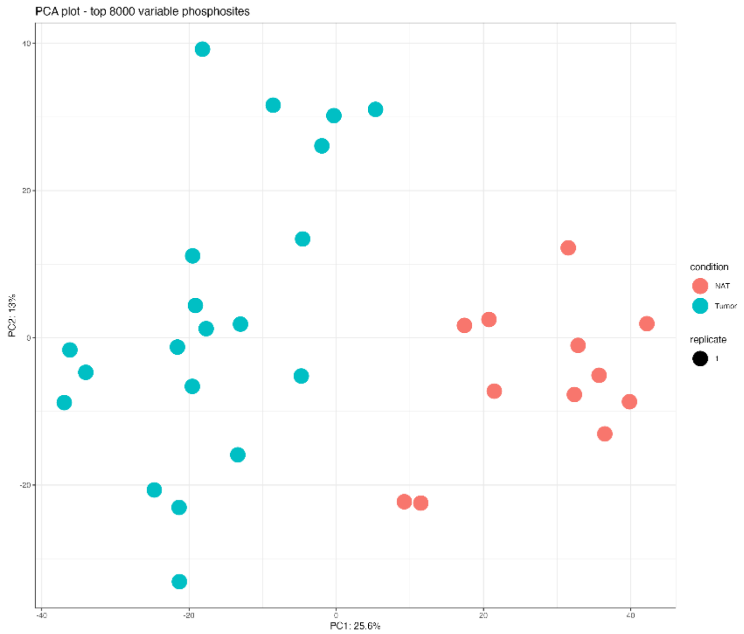


**Supplementary figure S13**. PCA plot of ccRCC phosphoproteomics dataset after normalization.
